# Stabilized marker gene identification and functional annotation from single-cell transcriptomic data

**DOI:** 10.1101/2024.08.21.608838

**Authors:** Sandesh Acharya, Pathum Kossinna, Qingrun Zhang, Jiami Guo

## Abstract

With the rapid emergence of single-cell transcriptomics datasets, reproducible marker genes and functional annotation of cell type or state is becoming increasingly important. Conventional methods that rely on differential gene expression (DEG) analysis lack both consistency across datasets and functional annotations of selected markers. Here, we present scSCOPE, an R- based platform that utilizes stabilized LASSO (Least Absolute Shrinkage and Selection Operator) feature selection, bootstrapped co-expression networks, and pathways enrichment to identify reproducible and functionally relevant marker genes and associated pathways for cell type identification and functional annotation in scRNAseq datasets. Using 8 scRNAseq datasets of immune cell types from human and mouse tissues generated by different sequencing technologies, we show that scSCOPE outperforms other popular methods by automatically identifying marker genes and pathways with the highest consistency across all datasets. We also demonstrate that scSCOPE’s gene co-expression and pathway analyses provide in-depth molecular insights into the functionality of identified marker genes. We anticipate that scSCOPE will greatly improve cell type/state annotation and accelerate the design of experimental validation and functional investigations on cell heterogeneity.

## Introduction

Single cell RNA sequencing (scRNAseq) enables high throughput profiling of transcriptomics for millions of cells at a time, and has transformed our understanding of cell heterogeneity, physiological state, and function in varied tissues across development and diseases^1–7^. Cell type identification in scRNAseq requires clustering of cells into distinct cell types based on their gene expression profiles, followed by the identification of marker genes associated with each cell type^8^. Currently, marker gene selection in scRNAseq data is an error-prone task. This is because common marker gene identification methods rely solely on differential gene expression (DEG) analysis, where the highest differentially expressed genes are often selected as cell-type specific markers^9^. Similarly, to infer cell-type specific functionality, current state-of-the-art methods use DEGs found in each cell type as inputs to look for pathway enrichments using databases such as KEGG, Gene Ontology^10–13^. Since gene expression can be sensitive to technical and biological variations in sample collection and sequencing platforms, these DEG-based methods suffer from instability across technical/biological variations and experimental conditions^33–35^. Further, since only the expression enrichment is considered, these selected markers often fail to offer cell-type specific functional annotations. To infer cell-type specific functionalities, pathway enrichment analysis of the DEGs will have to be performed separately. These pathways are then ranked based on enrichment scores or p-values, which are directly influenced by the number of DEG inputs. Various state-of-the-art methods including Mast, Deseq2, Bimod, Wilcox Rank Sum Test, Roc, DESingle have been developed to identify DEGs in scRNAseq datasets^14–20^. These DEG methods can be classified into two categories: a sample-level approach, which aggregates counts per cluster into pseudo bulks, and a cell-level approach, which models individual cells using generalized mixed effect models^21^. Both types of methods focus on mitigating the challenges associated with scRNAseq data’s inherent bimodality, dropout events, and technical variations^14–18,20,21^. However, there are two major limitations associated with these methods: (1) They analyze one gene at a time only based on expression values and do not consider gene-gene interactions, and therefore can be extremely sensitive to technical variations. Consequently, marker genes identified for the same cell type can differ across different studies, particularly for rarer cell types or transient cell states that are not well characterized; (2) Secondly, these techniques lack the capability to functionally annotate each marker gene in a particular cell type. As a result, researchers must manually choose a few marker genes from the pool of DEGs for experimental validation, based solely on their differential expression, and without functional insights.

Genes do not operate alone and hundreds of genes can be regulated by the same sets of transcriptional drivers. Incorporation of gene co-expression along with DEG analysis can help improve identification of marker genes and pathways that not only are enriched in each cell type of interest but also represent functionally important molecular signatures of cell types/states that are stable across datasets. The identification of such stable, cell type-specific molecular signatures is critical for generating accurate cell atlases and functional assessments. Currently, complex multi-gene models suffer from statistical instability, which results in inconsistencies when inputs are slightly altered^22–25^. This problem is particularly pronounced in scRNAseq analysis due to the variation associated with sequencing techniques and downstream data analysis^21^.

To overcome these limitations, we have developed scSCOPE (single-cell Stabilized COre gene and Pathway Election), which utilizes stabilized LASSO (Least Absolute Shrinkage and Selection Operator) feature selection, bootstrapped co-expression networks, and pathways enrichments to identify stable and functionally relevant marker genes and associated pathways for cell type identification and functional annotation using scRNAseq datasets^24–29^. scSCOPE is an extension of SCOPE, our previously established bulk RNAseq analysis method that empowers the synergy between co-expression analysis and regularized multiple regressions to provide stable and robust predictions of marker genes and pathways^28^.

We performed a systematic benchmarking of scSCOPE and other state-of-the-art methods^15–20^ across 8 scRNAseq datasets including 6 human PBMC (Peripheral Blood Mononuclear Cell)^30^ datasets and 2 mouse immune cell datasets generated by different sequencing technologies^31,32^. Our results show that scSCOPE accurately identifies marker genes and pathways that (i) show a high degree of gene co-expression, an indicative of involvement in cell-type specific functional programs; (ii) represent core cell-type specific molecular signatures that can be reliably captured regardless of technical variations across different scRNAseq datasets. Furthermore, scSCOPE automatically annotates each marker gene with enriched pathways and gene co-expression to facilitate cell-type specific functional annotations and validation.

## Results

### The Framework of scSCOPE

To develop scSCOPE from SCOPE, sparse regression and sparse correlation models were implemented to deal with sparse matrices associated with scRNAseq datasets. To improve stability, bootstraps on sub-sampled datasets were used when calculating the co-expression networks (in addition to SCOPE’s bootstraps on LASSO regression). Additionally, we implemented an integrated analysis of multiple modalities (regression, gene-gene co-expression, pathway enrichments and differential expression) to identify marker genes and pathways in single-cell RNA-seq data.

The input for scSCOPE is a clustered single-cell RNAseq dataset with an expression matrix (Fig. 1a). Based on scRNAseq data clustering (Fig. 1a), scSCOPE begins by running a bootstrapped logistic LASSO (see Methods) to identify “core genes” that robustly separate two groups of cells in multiple iterations^27^ (Fig. 1b). These “core genes” then undergo bootstrapped co-expression network analysis (Methods) to identify their stably co-expressed genes (“secondary genes”) (Fig. 1c)^25^. The “core-secondary” gene pairs are subsequently subjected to pathway enrichment analysis (Methods; Fig. 1d), leading to a collection of pathways enriched in each cell type (Fig. 1e)^29^. Next, the “core-secondary” gene pairs are ranked based on their pairwise correlations and enrichment in different pathways. Marker genes are then selected from all the genes in the top core-secondary gene pairs based on their differential expression (Methods, Fig. 1f, g). All the marker genes identified by scSCOPE are annotated with the top pathways they are associated with. This provides important functional annotations of the selected marker genes (Methods). As a final step, scSCOPE employs a unique ranking system to assess the identified pathways by integrating the impact of both gene expression and co-expression (Methods). Taken together, scSCOPE harnesses information from multiple modalities to identify highly stable and functionally informative cell-type specific marker genes and pathways using scRNAseq data.

**Fig. 1:**
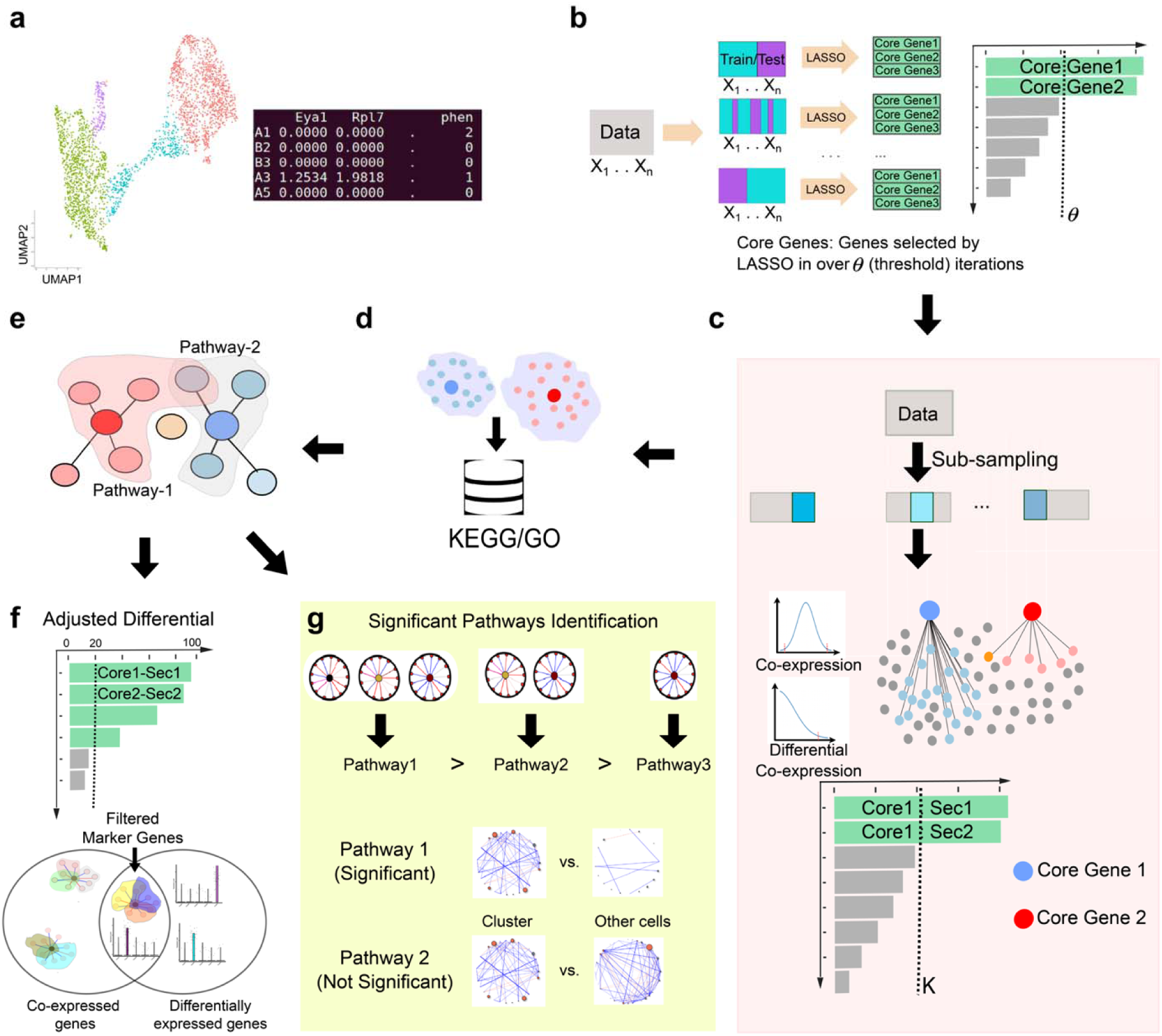
Overview of scSCOPE Framework. **a** scSCOPE requires a gene expression matrix along with cluster annotations as input. **b** The input data undergoes Iterative Sparse Lasso Regression to identify genes capable of segregating two distinct groups. In each iteration of LASSO regression, data is split into training and testing groups with the same cluster composition as in the original dataset. Only those genes consistently chosen in over a threshold (*θ*) of iterations are designated as Core Genes. **c** Core Genes are then subjected to Bootstrapped Co-expression analysis, where all the genes significantly co-expressed or differentially co-expressed with the core genes are identified as secondary genes. This analysis is run in a 60% sub-sample of the original data for 100 iterations and only stable gene interactions that appear in more than K iterations out of 100 are selected. **d** Each core gene, and its stable secondary genes identified through Co-expression analysis, then undergoes Pathway enrichment analysis. **e.** Pathway analysis can identify not only pathways enriched in the cluster but also core-secondary gene interactions and their involvement across multiple pathways. **f, g** The results from co-expression analysis, differential expression, and pathway enrichment are analyzed together to determine marker genes and pathways for the cluster of interest. Each of these steps are repeated and run in parallel for each analysis.

### scSCOPE identified marker genes that are stable across datasets and relevant to cell-type specific functions

By integrating bootstrapped re-sampling and regularized regression, co-expression networks, and pathway enrichment, scSCOPE will identify (i) marker genes that show the highest levels of gene co-expression that are potentially important for orchestrating cell-type specific functions and (ii) stable marker genes and pathways that are consistent across scRNAseq datasets.

To assess the accuracy, stability, and functional significance of the markers and pathways identified by scSCOPE, we collected 6 scRNAseq datasets on human PBMCs (Peripheral Blood Mononuclear Cells) generated from different sequencing technologies and 2 mouse immune cell datasets^30–32^. PBMCs, which include lymphocytes (T cells, B cells, and NK cells), monocytes, and dendritic cells, are one of the most well characterized cell types with identified cell markers and signaling drivers along each lineage^36^. The two mouse datasets are the GSE109999 and Tabula Muris datasets^31,32^. GSE109999 is a dataset of FACS-sorted immune cells (B-cells, T-cells, Granulocytes, Erythroblasts, and Progenitor cells) sequenced using CEL-seq2 technology^32^. As the cell types were determined prior to sequencing, this dataset is considered a gold-standard dataset (from here on referred to as “gsBlood” dataset). Tabula Muris is an extensive collection of single-cell transcriptome data obtained from around 100,000 cells representing 20 different organs in mice, including bone marrow (immune) cell types with pre-defined clusters that we analyzed here^31^.

To assess scSCOPE in comparison to alternative state-of-the-art methods including Deseq2, Wilcox, ROC, Bimod and MAST, we measured the performance of each method on their marker gene identification in all datasets. We first noted that scSCOPE identified a small number of marker genes (Fig. 2a) compared to all the DEGs provided by other methods (Supplementary table 1). DEGs were filtered based on their adjusted p-value (<0.05) and abs(logFC) (> 0.25). To compare the stability of marker gene identification, we used the 6 human PBMC datasets^30^. For DEGs identified by other methods, we chose the top 50 genes ranked by the highest average log2 fold change or lowest p-values to compare with scSCOPE. The stability of the top DEGs identified by each method and scSCOPE markers were then tested across the different datasets for each cluster (Fig. 2b). We found that scSCOPE markers showed the highest level of consistency across all 6 datasets compared to all other methods (Fig. 2c,d). This suggests that scSCOPE’s incorporation of gene co-expression networks improves the stability of marker gene identification as compared to conventional methods that are based on single-gene differential expression analysis.

**Fig. 2:**
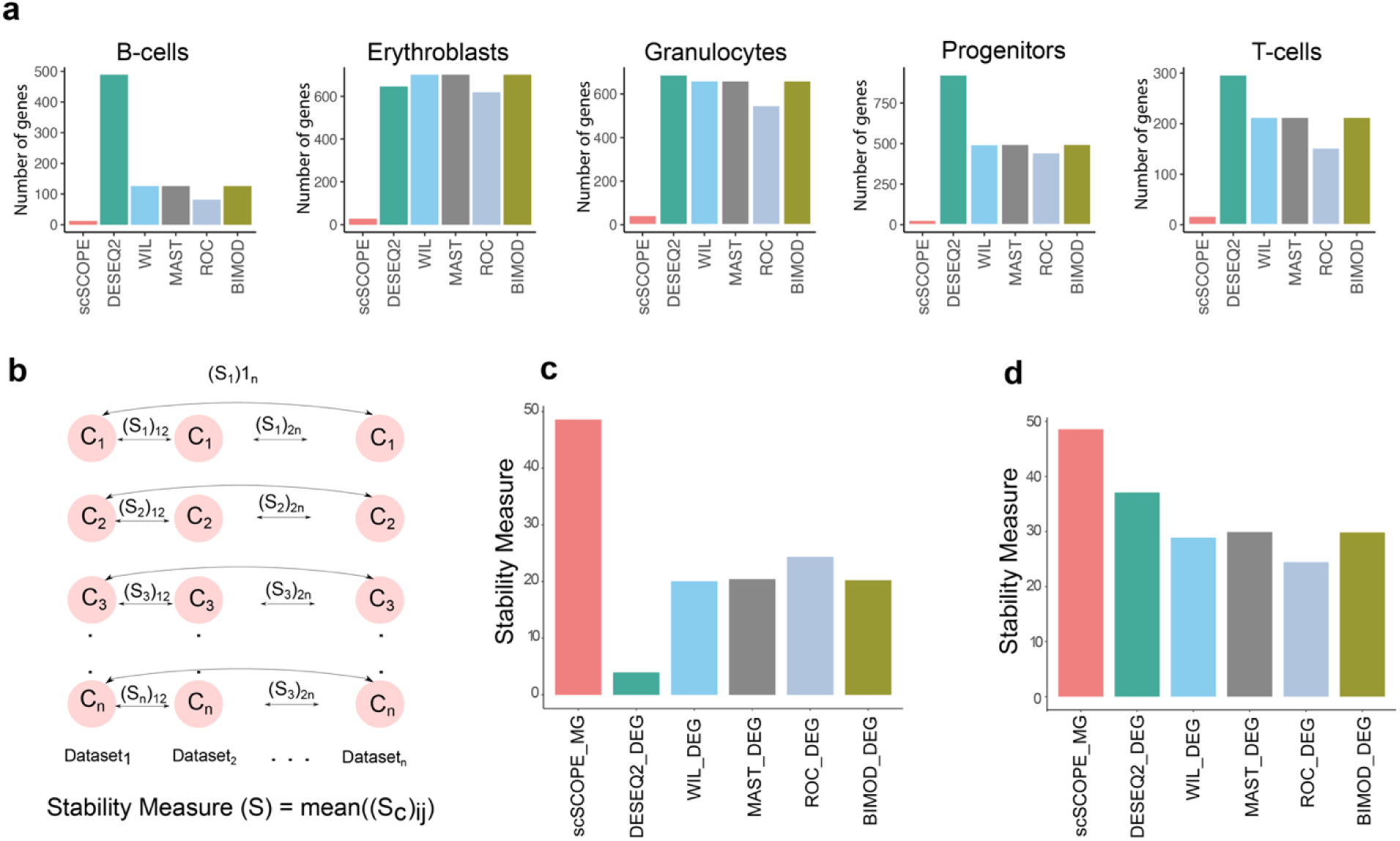
scSCOPE identifies fewer but more stable marker genes. **a** Bar Plots showing the number of marker genes identified by scSCOPE as compared to the DEGs identified by other methods. **b** The stability assessment of each method in identifying marker genes involves calculating the stability measure (S) based on the common marker genes detected within the same cluster across various datasets. This measure (S) is defined as the average of all (S_c_)_ij_ values, where (S_c_)_ij_ represents the percentage of common Marker Genes identified between dataset “i” and dataset “j” within cluster “c”. Here, “c” denotes the cluster of interest, “i” corresponds to dataset 1, and “j” corresponds to dataset 2. The (S_c_)_ij_ value is computed for all possible pairwise combinations of “i” and “j” for each cluster, and subsequently, these values are averaged to derive the overall stability measure (S). **c,d** Bar plots illustrating the stability measure of different methods for identifying marker genes in human PBMC datasets^1^. Top DEGs are selected based on their p-values in **c**, and the average log fold change in **d**.

Since scSCOPE considers the level of gene co-expression as a criterion for marker gene selection, we next assessed gene co-expression levels of scSCOPE-markers among all the DEGs using the Wilcox rank sum test. We first identified DEGs for each cell type in the gsBlood dataset, which were then subjected to “hub-gene” analyses using the String database and Cytohubba plugin within the Cytoscape application^37–39^. “Hub genes” are genes with the highest number of co-expressed genes^40^. We observed that the top hub genes consistently aligned with the scSCOPE-markers (Supplementary Fig. 1a-c). For example, 10 out of the top 15 hub genes in B-cells, 8 out of the top 10 hub genes in T-cells, and 9 out of the top 10 hub genes in Granulocytes were identified as scSCOPE-markers. This high degree of convergence suggests that scSCOPE indeed automatically identifies markers that show high levels of gene co-expression. The hub genes not selected as markers by scSCOPE are denied using criteria including pathway enrichment and differential expression.

### scSCOPE provides functional annotations to cell-type specific markers

scSCOPE’s implementation of gene co-expression and pathway enrichment for marker identification automatically provides functional annotations to the selected markers. All the marker genes and their associated pathways are listed in Supplementary Table 2. The gene co-expression networks across multiple pathways can be visualized by Gene Network Plots generated using the “geneNetwork” function from the R-implementation of scSCOPE. As an example, scSCOPE identified *Cd3d* as a marker for T-cells (Fig. 3a) and automatically revealed that *Cd3d* exhibits significant co-expression with genes in T- cell specific pathways, including Th17 cell differentiation, Th1 and Th2 cell differentiation, and T-cell receptor signaling pathways (Fig. 3b-e). In addition, the gene network plot for *Cd3d* in hematopoietic cell lineage pathway reveals that *Cd3d* is positively co-expressed with genes specific to T-cell speciation and differentiation, including *Cd2, Cd7, Cd3g, Cd3e, Itgb3, H2-Ab1*, while negatively co-expressed with non-T-cell specific genes like *Cd9, Kit, Itga4, Cr1l, Cd22* and *Cr2* (Fig. 3f, g). Thus, scSCOPE’s annotation is consistent with *Cd3d*’s known function in encoding a component of the T-cell receptor (TCR)/CD3 complex that drives T-cell activation and differentiation^41^.

**Fig. 3.**
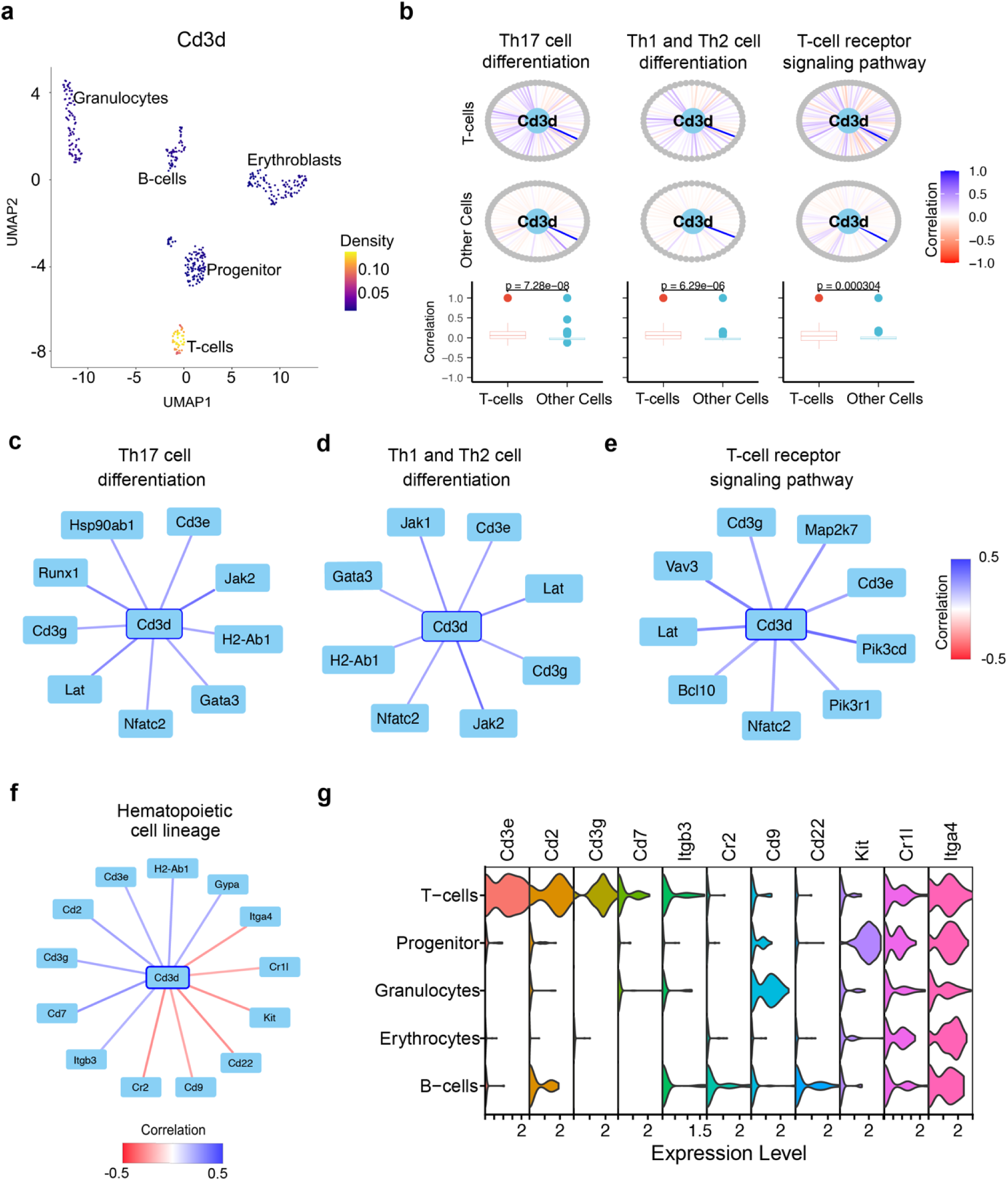
The role of *Cd3d* in T-cells as revealed by scSCOPE. **a** Density Plot showing the relative expression of *Cd3d* gene in different cell types in the gsBlood dataset. **b** The gene network plots for *Cd3d* gene provide a visual representation of its extensive interactions with genes across diverse pathways. Within each plot, surrounding *Cd3d* are gray circles representing all other genes within the pathway. The lines connecting *Cd3d* to these genes indicate Pearson’s correlation coefficient, ranging from −1 (red) to +1 (blue), reflecting the strength and direction of correlation. Correlations are separately calculated for two distinct groups: in this case T-cells and all other cell types. Boxplots accompanying the plots contrast the distributions of correlations between groups. *p*-values from the Kolmogorov-Smirnov test are provided to assess the statistical significance, with the null hypothesis stating that two samples are drawn from the same distribution. **c,d,e,f** Gene Network Plots are zoomed in to visualize only the top genes which show highest correlation (|Correlation| ≥ 0.2) with *Cd3d* across different pathways in T-cells. Lines are colored according to the Pearson’s correlation between two genes from −0.5 (red) to + 0.5(blue). **g** Violin Plots for genes significantly correlated with *Cd3d* in the Hematopoietic cell lineage pathway.

The gene network plot generated by scSCOPE is particularly useful to facilitate target gene prediction when a transcription factor is identified as a marker gene. For example, scSCOPE identified *Tcf7* (transcription factor 7), a gene encoding the transcription factor Tcf1 as a marker gene for T-cells in the gsBlood dataset (Supplementary Fig. 2a). In total, *Tcf7* gene exhibited significant co-expression (absolute value ≥ 0.2) with more than 1700 genes across the entire dataset. scSCOPE refined these gene co-expression to focus only on T-cell specific pathways, thereby isolating and prioritizing the most relevant gene interactions for T-cells. Gene network plots show that *Tcf7* is significantly co-expressed with 57 genes among T-cell related pathways (e.g. T-cell receptor signaling pathway, Th17 cell differentiation, Th1 and Th2 cell differentiation pathways) (Supplementary Fig. 2b). Notably, 49 out of these 57 genes have binding sites for Tcf1 in their promotor regions^42,43^, and 38 out of these 49 genes have been validated using ChIPseq experiments^43^ (Supplementary Fig. 2c). Of the remaining eight genes, four (*Maml2, Irf4, Hras, and H2-DMa*) are regulated by Runx1, a target gene of Tcf1^42^.

Collectively, these results demonstrated that scSCOPE’s gene co-expression and pathway analyses provide enriched and in-depth molecular insights into the functionality of identified marker genes in each cell type.

### scSCOPE identified a new B-cell marker *Il7r*

scSCOPE utilizes a multi-step filtering process to pinpoint informative marker genes for cell type classification in scRNAseq data. Genes that can distinguish the cell-type of interest, show extensive co-expression with genes enriched in cell-type specific pathways, and that are differentially expressed are selected as markers in scSCOPE (Supplementary Fig. 3a). As a result, genes that rank low in DEG analysis but excel in co-expression and pathway analysis may be identified as markers by scSCOPE.

As an example, scSCOPE identified *Il7r*, a gene with a low differential expression score but substantial co-expression, as a marker gene for B-cells in the gsBlood dataset (Supplementary Table 2). Based on expression enrichment, Il7r ranks low among DEGs for B-cells (MAST: 46th, DESeq2: not identified as a DEG, Wilcox: 46th, ROC: 46th, Bimod: 46th) and is consequently not selected as a marker from previous scRNAseq studies (Supplementary Fig. 3b). However, when considering both the strength of gene co-expression and involvement in pathways, *Il7r* exhibits strong correlations with three core genes identified for B-cells (Supplementary Fig. 3c) and is enriched in two pathways specific to B-cell functions and is therefore identified as a B-cell marker by scSCOPE. In contrast, while genes such as *Cd74* and *Igkm* rank high in DEG analyses in B-cells, they do not exhibit significant co-expression or pathway enrichment and are consequently disregarded by scSCOPE as markers (Supplementary Fig. 3b,c).

Due to its higher degree of co-expression and involvement in multiple pathways, it ranks higher in the “adjusted differential” metric compared to other genes like *Cd74* and *Igkc*, which have higher fold changes but lower degrees of correlation and pathway involvement.

Il7r is a cell membrane receptor for the interleukin-7 protein^44–47^. Extensive literature has reported the critical role of Il-7R signaling in the survival and maintenance of lymphocytes^44–51^. In the context of B-cells, IL-7R signaling has a well-defined role in promoting the proliferation and survival of B-cell progenitors^45,47–49,51^. To demonstrate the accuracy and depth of scSCOPE’s functional annotation of marker genes, we next evaluated the scSCOPE generated *Il7r* annotation alongside the known functions of IL- 7R signaling in B-cells. We employed scSCOPE on the Tabula Muris dataset focusing specifically on the bone marrow. This dataset incorporates data from two experiments with distinct methodologies for cell isolation: droplet technology and FACS sorting. The droplet dataset offers broader coverage across B-cell stages, while the FACS dataset provides higher depth sequencing. The two datasets encompass diverse and partially overlapping developmental stages of B-cells. We leveraged both datasets to include all B-cell subtypes in our analysis.

The development of B-cells is dependent on the sequential DNA rearrangement of the immunoglobulin loci that encode subunits of the B cell receptor^47^. The hematopoietic progenitor cells undergo the B-cell lineage specification and commitment process from Pre-Pro, Pro-B (further divided to Early-Pro and Late-Pro), Pre-B, immature-B, to naïve B-cells^50,51^. Cell proliferation and survival, two major events during B-cell development, are both known to be regulated by IL-7R signaling, especially during the Pro-B and Pre- B stages^47–49^. Importantly, scSCOPE accurately identified *Il7r* as a marker specifically for Late-Pro and Pre-B stages.

Expression wise, consistent with its marker status, *Il7r* indeed shows the highest expression levels at the Late-Pro and Pre-B stages compared to all other stages (Fig. 4a, b). scSCOPE’s Gene Network Plots highlighted that *Il7r* is significantly co- expressed with genes enriched in cell cycle and survival-related pathways (Fig. 4c, f). For cell cycle regulation, scSCOPE revealed that *Il7r*’s shows an increased co- expression correlation in the cell cycle pathway at Late-Pro compared to Early-Pro stage (Fig. 4c). Indeed, *Il7r* is positively co-expressed with many cell-cycle related genes like *Bub1*, *Mki67*, *Cd72*, *Cdc20*, *Ndc80*, *Top2a*, *Cdc25b* that promote B-cell proliferation in the Late-Pro stage compared to Early-Pro and Pre-B stages (Fig. 4d, e). For cell survival regulation, IL-7R signaling is known to activate downstream STAT5 transcription factor and PI3K-AKT signaling to promote cell survival along B-cell development^47^. Consistently, scSCOPE’s co-expression and pathway enrichment analyses revealed that *Il7r* is co-expressed with genes enriched in JAK-STAT, PI3K- AKT, and related pathways such as the FoxO pathway that is repressed during cell survival, and the Cytokine-Cytokine receptor interaction pathway important for B-cell differentiation (Fig. 4f). Interestingly, from the Late-Pro to Immature B cell transition, *Il7r*’s co-expression patterns in these pathways show a shift from a predominantly negative correlation to positive correlation (Fig. 4f), indicating that IL-7R signaling activity is dynamically regulated during this process, consistent with experimental findings^47–49^. Although *Il7r* gene expression or co-expression correlation is not a direct readout of IL-7R signaling activity, these analyses demonstrates scSCOPE’s great sensitivity and accuracy in providing insights into the dynamic regulation of *Il7r* and related signaling pathways during early B-cell differentiation.

**Fig. 4:**
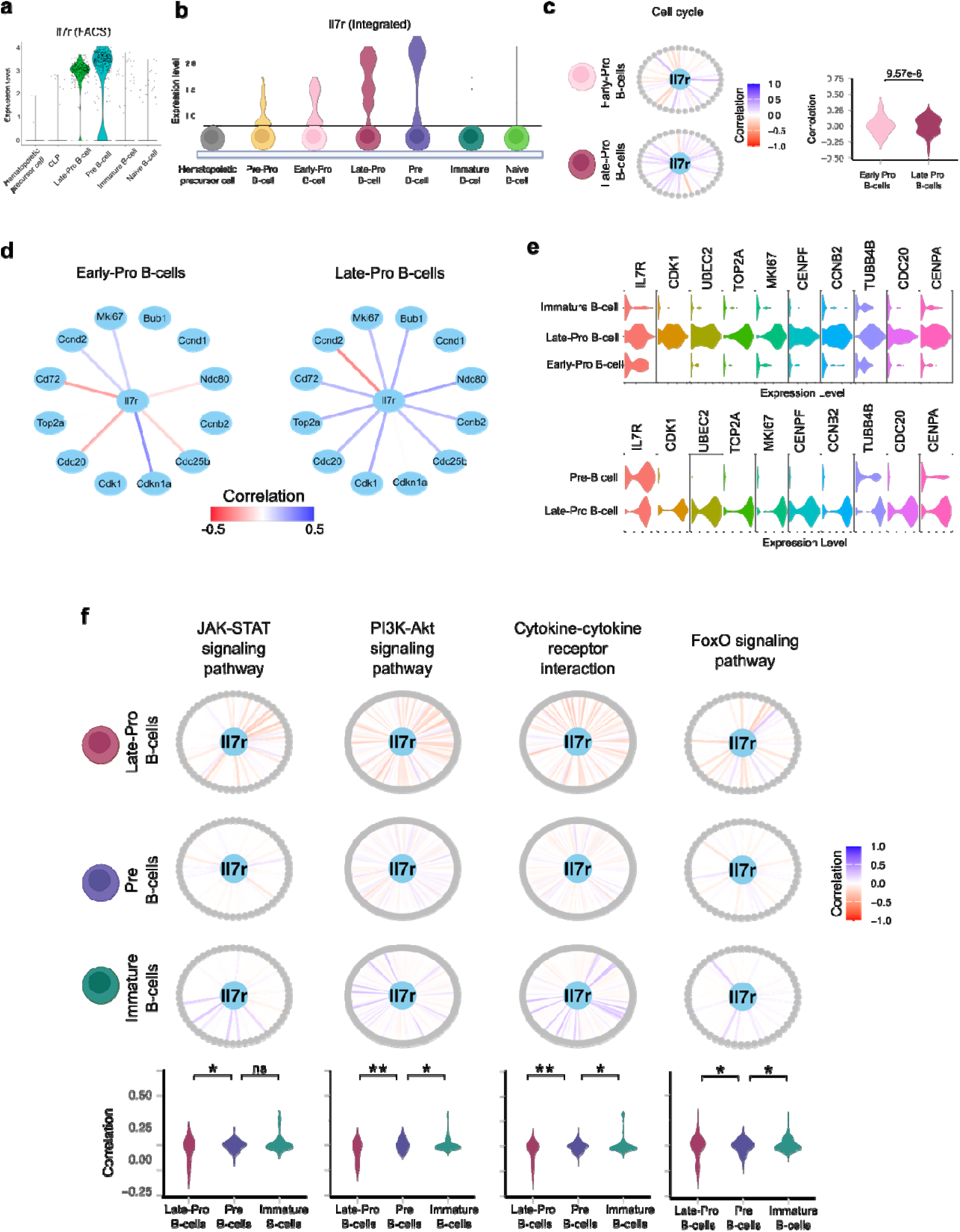
scSCOPE identified *Il7r* gene as a marker gene for Late-Pro and Pre-B cells. **a** Violin Plots showing the relative expression of *Il7r* gene across different B-cell stages in FACS dataset of Tabula Muris. **b** Relative expression of *Il7r* gene in different B-cell stages generated by integrating the FACS and Droplet datasets from Tabula Muris. **c** Gene Network Plots for *Il7r* in Cell Cycle Pathway in Early-Pro and Late-Pro B-cells of the Droplet-Tabula Muris dataset. *Il7r* gene is placed in the middle with all other genes expressed in the pathway placed in the periphery. The lines connecting *Il7r* to these genes indicate Pearson’s correlation coefficient, ranging from −1 (red) to +1 (blue), reflecting the strength and direction of correlation. Correlations are separately calculated for Early-Pro B-cells and Late-Pro B-cells. Violin Plots accompanying the plots contrast the distributions of correlations between these groups. *p*-value from the Kolmogorov-Smirnov test is provided to assess the statistical significance, with the null hypothesis stating that two samples are drawn from the same distribution. **d** Correlation Plots for *Il7r* with selected genes in the Cell Cycle Pathway across Early-Pro and Late-Pro B-cells. Lines are colored according to the Pearson’s correlation between two genes from −0.5 (red) to + 0.5(blue). **e** Violin Plots show the relative expression of G2M- phase genes in different clusters in the FACS and Droplet Tabula Muris datasets. **f** Gene Network Plots for *Il7r* gene in Late-Pro, Pre-B and Immature B-cells of the FACS- Tabula Muris dataset. Correlations are separately calculated for three groups. Violin Plots accompanying the plots contrast the distributions of correlations between these groups. P-values (Kolmogorov-Smirnov test) are denoted: *, p < 0.05; **, p < 0.01; ***, p < 0.0001; ns, p ≥ 0.05. Corresponding p-values for Late-Pro *vs.* Pre-B cells are: 0.0023 (JAK-STAT), 0.00013 (PI3K-Akt), 0.00034 (Cytokine-cytokine), 0.010 (FoxO).

Corresponding p-values for Pre-B *vs.* Immature B-cells are: 0.054 (JAK-STAT), 0.024 (PI3K-Akt), 0.047 (Cytokine-cytokine), 0.016 (FoxO).

### scSCOPE enabled functional prediction of the unannotated gene Gm8292

In cases where scSCOPE identifies unannotated or poorly characterized genes as markers, their Gene Network Plots can offer valuable clues about their potential functions within the cell type of interest. As an example, scSCOPE identified *Gm8292* as a marker gene for hemopoietic progenitor cells in the gsBlood dataset (Fig. 5a). *Gm8292* (ENSMUSG00000100215) is a mouse pseudo-gene on chromosome 1. Its expression has been associated with conditions such as cholestatic intestinal injury and TCDD-induced cleft palate^52,53^. However, its function remains unknown. scSCOPE’s Gene Network Plot of *Gm8292* revealed its significant correlation with multiple genes in the “Cell Cycle” pathway (Fig. 5b). Interestingly, *Gm8292* showed a negative correlation with many genes involved in “Cell cycle arrest” including *Ticcr*, *Cdkn2d*, *Chek2*, *Mad2l1*, *E2f2*, *Cdkn2c*, and *Pkmyt1* (Fig. 5c). Hence, via scSCOPE, we predict that the newly identified hemopoietic progenitor cell marker *Gm8292* is involved in cell cycle regulation, and likely plays a role in the promoting cell cycle progression.

**Fig. 5:**
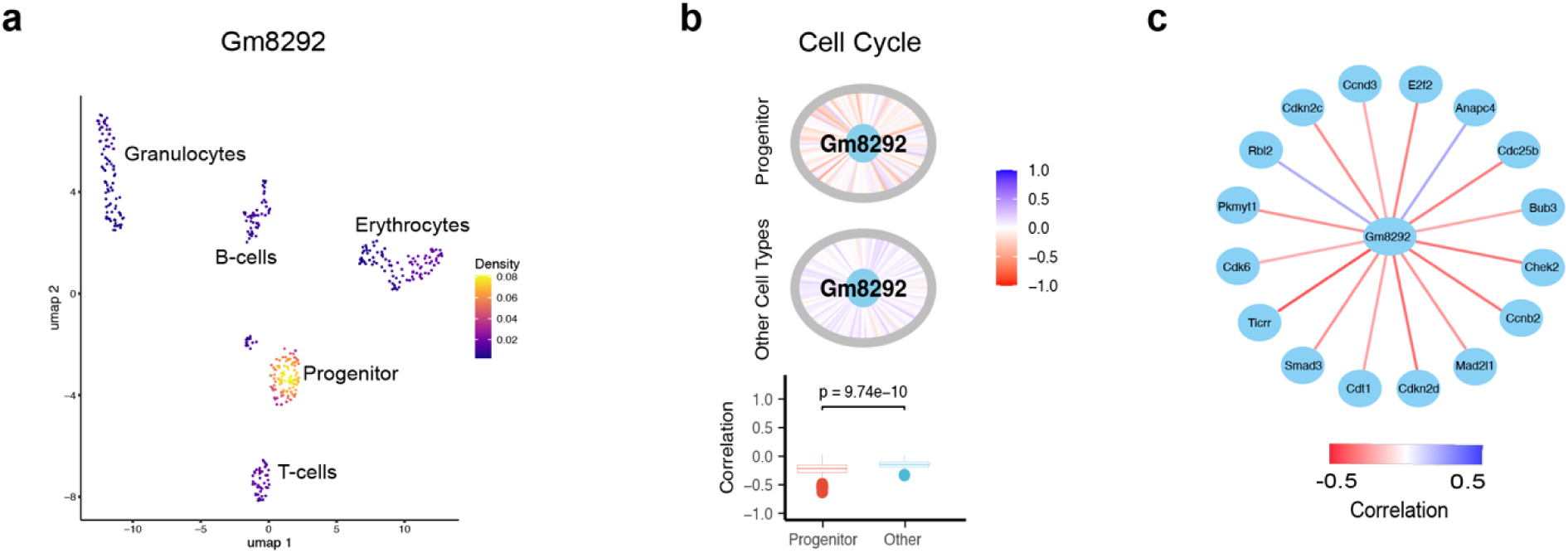
scSCOPE’s functional annotation of an un-annotated gene *Gm8292*. **a** Density Plot showing the expression level of *Gm8292* in different clusters of gsBlood dataset. **b** Gene Network Plot for *Gm8282* shows that this gene is co-expressed with many genes within the Cell Cycle pathway. All the genes in Cell Cycle pathway are placed in the circumference of the circle (Grey color) and *Gm8292* gene is placed in the center (Light blue). Lines represent the strength and direction of Pearson’s correlations between *Gm8292* and all other genes from −1 (red) to +1 (blue). Boxplots contrast the distributions of two sets of correlation in the Progenitor cells vs all other cell types. The Kolmogorov-Smirnov test was performed, with the null hypothesis being that two samples are drawn from the same distribution. **c** Gene Network Plots showing correlations between *Gm8292* and selected genes from the Cell Cycle Pathway (with absolute correlation > 0.2).

### scSCOPE identified more cell-type enriched pathways and more genes within each pathway

As many genes are co-regulated within the same signaling pathways, incorporation of co-expressed genes along with the top differentially expressed genes will help identify more pathways that are significantly enriched in a cell type of interest. However, conventional co-expression analysis with an extensive list of DEGs can significantly escalate computational complexity and time. scSCOPE’s pathway enrichment method uses both core genes (identified by bootstrapped LASSO) and secondary genes (co- expressed with core genes) as inputs to identify enriched pathways in each cluster of interest. Compared to common pathway analyses such as GSEA and ORA^29,54^ that use DEGs to find enriched pathways, scSCOPE exceled by identifying (i) a higher number of enriched pathways and (ii) more stable pathways across different datasets (Supplementary Fig. 4a, b, Supplementary Table 1, 3). For example, in the gsBlood dataset, pathways such as the “IL-17 signaling pathway” which is important for granulocytes development was not identified by ORA or GSEA for Granulocytes but was identified by scSCOPE.

To highlight the relevance of enriched pathways in each cell type, scSCOPE employs a unique ranking system to indicate the significance of the identified pathways. scSCOPE calculates the weights of each pathway to generate “CorrExpress” values (see methods) by incorporating the co-expression correlation and expression differences of all genes within a pathway (Fig. 1a, Methods). The top pathways for each analysis are then ranked based on the absolute values of CorrExpress, referred to as “Final Measure”. As a result, scSCOPE outputs a Pathway Bar Plot to identify the top pathways relevant for a cell-type as well as Pathway Network Plots to study the patterns of expression and co-expression of genes in the pathway in different cell types (Supplementary Fig. 4c, d). Pathway Network Plots can be generated using the “pathwayNetwork” function and Pathway Bar Plots can be generated using the “pathwayBar” function from R-implementation of scSCOPE.

In addition to pathway identification, scSCOPE also outperformed other methods in the number of genes it identified within each pathway, for all the pathways enriched in each cell type (Supplementary Fig. 5, Supplementary Table 4). For instance, scSCOPE identified a total of 28 genes in the B-cell receptor signaling pathway, while other methods identified fewer than 16 genes for the same pathway. This expanded gene set within each pathway not only substantiates the robustness of pathway enrichment analysis, but also provides researchers with a more comprehensive list of potential regulators and effectors within these pathways.

## Discussion

With the advancement of single cell transcriptomics techniques, a vast amount of single-cell transcriptomic profiling data has been generated from humans and other species^31^. While these information-rich datasets enable researchers to readily capture all cell types and states from complex tissues, they also engender challenges for consistent identification of cell type-specific markers and pathways that reliably inform the core cell molecular features of a specific cell type.

Conventional methods perform marker gene identification entirely based on differential expression of genes (DEGs). Although DEG methods analyze the “enrichment” of genes to distinguish cell types, their focus on the expression of singular genes can make them sensitive to sample collection and technical variations, resulting in instability across datasets and a lack of functional annotations of the selected markers or cell types. This issue becomes more prominent in rarer cell types or states that are lower in abundance or sensitive to environmental stimuli. This lack of consensus in marker identification and functional annotation of the newly identified cell types or cell states presents a major challenge for further investigation.

Here, we present scSCOPE, an optimised toolbox for single-cell RNA-seq based cell- type functional annotation. To the best of our knowledge, scSCOPE is currently the only computational tool that implements gene co-expression, pathway enrichment and differential expression to identify marker genes in single cell transcriptomics data.

scSCOPE is also the first tool to use genes co-expression to identify and rank pathways in single-cell transcriptomics data. To promote its application, scSCOPE is implemented as an open-source R-implemented tool (https://github.com/QingrunZhangLab/scSCOPE) to enable fully automated scRNAseq based cell-type functional annotation.

In comparison with other computational approaches that require manual inference for marker gene selection and pathway enrichment analysis based on differential expression, scSCOPE enables automatic identification of markers that are not only cell- type specifically enriched but also highly interactive in cell-type specific pathways in an unsupervised manner. Further, using 8 scRNAseq datasets of well-characterized immune cell types in humans and mice by different sequencing technologies (i.e., SMART-seq2, 10X_v2, 10X_v3, Dropseq, CelSeq, inDrops), we benchmarked scSCOPE against other state-of-the-art methods (DESeq2, Wilcox Rank Sum, MAST, ROC, Bimod). Overall, our results demonstrated that scSCOPE (i) showed the highest degree of stability in cell-type specific marker gene and pathway identification across all datasets ; (ii) was able to identify not only the well-established marker genes but also new marker genes based on their extensive gene co-expression within the cell-type specific pathways, despite their relatively low expression enrichment; (iii) enabled the functional prediction of an unannotated marker gene, and (iv) lastly, identified more cell- type specific pathways and more enriched genes within each pathway. We anticipate that with these powerful advancements, scSCOPE will greatly improve cell type/state annotation and accelerate the design of experimental validation and functional investigations on cell diversity, particularly when it comes to rare cell types or transient cell states that are poorly characterized, highly dynamic, and sensitive to external stimuli.

As an example, we demonstrated that scSCOPE identified *Il7r* as an important marker gene for Late-Pro and Pre-B cells. Extensive experimental research^44,47,55–57^ has established *Il7r* as a well-known key gene for B-cell differentiation. Nevertheless, due to its relatively low differential expression score, no scRNAseq analysis methods, except for scSCOPE, identified *Il7r* as a marker gene. Importantly, while genes such as *Il7r* that encode surface membrane proteins (e.g., signaling receptors) can have powerful impacts on cell differentiation or cell state transitions, they often do not express at high levels and therefore are rarely identified as markers in scRNAseq datasets. The fact that scSCOPE selects *Il7r* as a marker gene highlights its unique ability to identify functionally key genes as cell type markers. Notably, Late-Pro and Pre-B cells represent transient cell states, transitioning from progenitors to immature B-cells. The scSCOPE’s identification of *Il7r* as a marker during these states highlights its high sensitivity and accuracy in capturing the gene signatures that potentially drive cell lineage/state transitions.

Since scSCOPE includes co-expressed genes in pathway enrichment, it identifies more pathways than other methods. Incorporation of co-expressed genes during pathway enrichment also adds a layer of consistency across different datasets. In addition, scSCOPE introduces a novel validation strategy for inferred pathways, based on the expression and co-expression patterns of genes. This innovative approach adds a layer of rigor to the analysis, improving the distinction between biologically meaningful pathways and potential artifacts. When many genes in a pathway exhibit both differential expression and differential co-expression in the cell type of interest, it indicates a potential significance of the pathway in driving cell-type specific biology.

Since the input for scSCOPE is a gene expression matrix of all the cells and the phenotype/cluster annotation for each cell, one limitation of scSCOPE is that it relies on accurate clustering. If the initial clustering is wrong, scSCOPE fails to identify unique marker genes and pathways for the cluster of interest. Due to the lack of solutions on simulating scRNA-seq data with a proper correlation structure consistent with genuine pathways, our current study does not include simulation assessments. The gene network analysis in scSCOPE provides more stability than conventional DEG methods, but still relies on good sequencing depth and coverage of the transcriptome. With continuing advancements in single-cell sequencing technologies, the accuracy and sensitivity of scSCOPE is expected to increase accordingly.

In conclusion, scSCOPE’s consideration of gene networks, novel pathway validation strategy, and comprehensive pathway enrichment analysis collectively not only refines our ability to identify critical genes and pathways specific to each cell type but also offers more biologically meaningful perspectives on cellular heterogeneity.

## Methods

### scSCOPE Framework

The SCOPE framework was designed to identify candidate genes and pathways separating normal and diseased tissues in bulk RNA-seq datasets. We have further improved the SCOPE framework to be applied directly to single cell datasets to identify cell-type specific marker genes and pathways.

### SCOPE-Stabilized LASSO selection

The initial step in scSCOPE involves the deployment of the LASSO algorithm to discern core genes capable of distinguishing between two groups^24^. Addressing the inherent instability associated with the LASSO algorithm, the SCOPE methodology uses bootstrapped LASSO regression to select genes exhibiting consistent behavior across multiple iterations^28,58,59^. However, the SCOPE methodology, which was developed for binary phenotypes and bulk RNA-seq data, could not be applied directly to single-cell data with multiple phenotypes. We therefore introduced “1 vs all” and “1 vs 1” logistic regression models to identify core genes associated with each cluster and differentiating between two clusters, respectively, in scSCOPE. Consistent with SCOPE’s methodology, LASSO regression iterates 200 times on sub-sampled data (split 70-30) and genes selected in over θ runs, termed Core Genes, are chosen for subsequent analysis. The default value for θ is 160 (80% of total iterations). In cases where no core genes are identified, the algorithm automatically selects the top five genes which appear most frequently in the LASSO iterations as core genes. To address the sparsity inherent in single-cell data, we used sparse matrices during LASSO regression.

### Co-expression and pathway analysis

Genes function within complex networks, interacting with other genes across various pathways to shape specific phenotypic traits^1,2,34^. Despite the good predictive performance of LASSO, it suffers from unstable selections of correlated variables and inconsistent selections of linearly dependent variables^28,58,59^. The original SCOPE framework utilizes Co-expression and Differential Co-expression analyses to reveal genes that are strongly associated with each core gene, which could be missed by LASSO feature selection^28^.

To quantitatively analyze these relationships, we computed pairwise correlations between core genes and all other genes using the corSparse function from the qlcMatrix package^60^, which accommodates for sparse matrices in single-cell data. Only those gene pairs that surpassed predefined thresholds for either differential co-expression or co-expression were deemed significant. These thresholds were meticulously determined. For correlation, we extracted the 97.5th percentile from a null distribution of correlations calculated among 1000 random genes in 100 rounds. We repeated this for both positive and negative threshold calculations. Similarly, for differential co-expression, we identified the 97.5th percentile from a null distribution of differential co- expressions (Correlation_group1_ – Correlation_group2_) between 1000 random genes across 100 rounds. Importantly, these threshold values were established separately for each cluster or analysis.

Only those genes demonstrating either pronounced co-expression or significant differential co-expression with the core genes, called secondary genes, were advanced for further analysis. To ascertain the stability of core-gene/secondary-gene correlations, we performed 100 bootstraps of co-expression analysis on a sub-sample, representing 60% of the entire dataset. Sampling was done on each iteration. This sub-sample preserved the original dataset’s cluster distribution. Only core-secondary gene pairs deemed significant in over K sub-sampled rounds were retained for further analysis. The default value for K was 80.

For each core gene and its associated secondary genes, we conducted an Over- Representation Analysis (ORA) as well as Gene Set Enrichment Analysis (GSEA) using the KEGG Pathway and Gene Ontology Database^10–12^. This analysis, executed through the WebGestaltR platform, applied a stringent false discovery rate (FDR) threshold of 0.05 to highlight pathways of notable significance^13^.

### Marker Gene Identification

After identifying gene pairs that exhibit notable correlation or differential co-expression between core and secondary genes, we introduced a metric known as “adjusted differential correlation” to effectively rank these pairs.

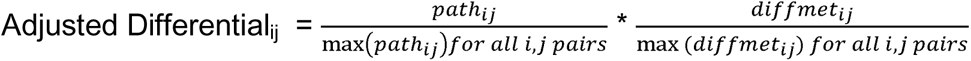, where

*i* = core gene, *j* = secondary gene, *path_ij_* = number of pathways the gene pair (*i, j*) is involved in, *diffmet_ij_* = max(*correlation_ij_*, *group1_correlation_ij_*, *group2_correlation_ij_*, *differential_correlation_ij_*) is the maximum of correlation of the two genes or differential correlation of the two genes between two groups.

The computation of adjusted differential correlation involves two components: the path ratio and diffmet ratio. The path ratio is the ratio of the number of pathways the core- secondary pair is enriched in to the maximum number of pathways any core-secondary pair is enriched in. Similarly, diffmet ratio is the ratio of diffmet values for the core- secondary pair to the maximum value of diffmet for the cluster. Thus, the adjusted differential metric considers both the degree of differential co-expression/correlation and the pathways associated with the gene pairs. In simpler terms, gene pairs that are linked to multiple pathways and display substantial correlation or differential co-expression hold more significance as compared to others.

To refine our focus, we established a threshold for adjusted differential correlation. Only gene pairs surpassing 20% of the maximum adjusted differential correlation within the cluster were deemed significant. In cases where the counts of identified marker genes are exceptionally low, this threshold can be relaxed to capture additional genes of interest.

Finally, we evaluated the fold change for each gene from the above pairs. Marker genes were then filtered using a threshold for both fold change (0.5) and the fraction of cells expressing the gene (0.45). This stringent criterion helped to pinpoint genes that play a substantial role in characterizing the specific cluster under analysis.

### Stability Calculation

We used human blood cell datasets from different platforms (10x-v3, 10x-v2, Dropseq, CELSeq, Seqwell and inDrops) generated by Ding et. al^30^ to test for the stability in DEG and pathways identified by different methods. To do this, we identified significant DEGs (adjusted p-value < 0.05) in each cluster of all the datasets using different methods (Wilcox Rank Sum, Bimodal analysis, MAST, DESEQ2, and ROC)^15–18,20^. We selected the top 50 DEGs (ranked by either their absolute average fold change or by their p-values) as marker genes from different methods. Next, we calculated overlaps between DEGs in each cluster of one dataset with the same cluster from another dataset. Pairwise overlaps over all clusters and all datasets were averaged to generate a new value “Stability Measure”. Similarly, we also calculated the Stability measure for marker genes identified by scSCOPE. We also calculated stability measures for the pathways identified by scSCOPE and other methods in the same datasets.

### Stable Pathways Identification

We performed pathway enrichment for each core gene and its surrounding genes for each cluster separately. For each identified pathway, we calculated the number of core- secondary gene pairs enriched in the pathway. Next, we calculated the CorrExpress measure to differentiate between the expression and correlation pattern in the cluster of interest and all other cells. CorrExpress measure was calculated separately for genes expressing positive differences (posCorrExpress) between two clusters and genes expressing negative differences (negCorrExpress) between two clusters.

For genes *i* with { (*exp_i_*)*_cluster_* - (*exp_i_*)*_othercells_*} ≥ 0:

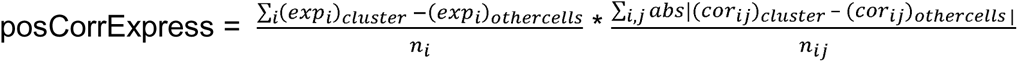

and for genes *i* with { (*exp_i_*)*_cluster_* - (*exp_i_*)*_othercells_*} <0:

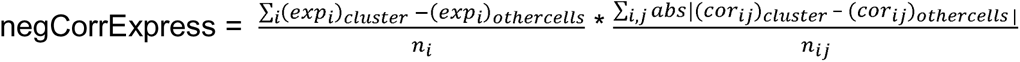

where ∑*_i_* denotes the sum over all genes in the pathway, (*exp_i_*)*_group1_* represents the average expression of gene *i* in the first group, (*exp_i_*)*_group2_* represents the average expression of gene *i* in the second group, and *n_i_* is the total number of genes in the pathway. Similarly, ∑*_i,j_* denotes the sum over all pairs of genes in the pathway, (*cor_ij_*)*_group1_* represents the Pearson’s correlation between genes *i* and *j* in the first group, (*cor_ij_*)*_group2_* represents the Pearson’s correlation between genes *i* and *j* in the second group, and *n_i_*, *j* is the total number of gene pairs in the pathway.

The calculation of the pathway difference metric involves two key steps. First, the differences in the expression levels of individual genes within the specified pathway between two groups are added together and normalized. This is done separately for genes expressing positive differences as well as genes expressing negative differences between two groups. Second, the absolute differences in pairwise correlations among all gene pairs across the two groups are calculated and averaged. These two resulting values are multiplied, creating a composite metric, “Final Measure”, that effectively combines the influences of both gene expression variations and correlation dynamics within the pathway. Finally, top pathways for each analysis are ranked based on the absolute values of posCorrExpress or negCorrExpress metrics. Moreover, scSCOPE extends its analysis by attributing significance levels to both expression and co-expression differences between two groups. This is achieved by randomly selecting an equivalent number of genes as those in the pathway and computing the expression and co-expression differences between the two groups. The resulting distributions for expression and co-expression differences from these random genes are then compared to those of genes from the pathway. This comparison is carried out using a Kolmogorov-Smirnov Test (K-S test), yielding separate p-values for both expression and co- expression differences^61^. The correlation and expression difference between two groups in each pathway can be visualized using Pathway Network Plots generated using visNetwork library in R^62^.

By integrating both gene expression and correlation aspects, the pathway difference metric offered a thorough evaluation of the pathway’s significance. This comprehensive assessment facilitated the prioritization and ranking of relevant pathways within each cluster.

### Accounting for Uneven datasets

Correlation analysis can be highly affected by the number of observations in each cluster of interest. We have incorporated a maximum sampling strategy to account for this problem. A threshold for maximum number of samples in each phenotype is used such that all different clusters have similar number of cells. This is implemented on each iteration of Logistic LASSO and Correlation analysis.

### Pathway Annotation for each marker gene and Gene Network Plots

To annotate each marker gene with pathways, we counted the number of genes in the pathway which were significantly correlated with our marker gene of interest. Those pathways with a higher degree of association with the marker gene were determined to be more significant for the marker gene in the cluster of interest. Gene Network Plots were generated based on Pearson’s correlation values between the marker gene and all other genes in the pathway across two groups. To look at the significance of correlation difference between two groups, the KS test was performed between pairwise correlations of group one with group two and was represented by a box plot /violin plot in the gene network plots. In cases where there were fewer than 100 gene-gene correlation values, sampling with replacement was carried out to take at least 100 correlation values for analysis.

### Hub Gene Identification using Cytoscape

A list of differentially expressed genes were input into the Cytoscape application and the full STRING network for the list was generated using the STRING database^37–39^. Next, we calculated node scores for each gene using the Cytohubba plugin and ranked them based on their degrees (interaction). The top 15 nodes/genes were selected as hub genes.

### Implementation in real datasets

We implemented scSCOPE in six immune cell datasets to validate the accuracy and applicability. Immune cell datasets were chosen because they were highly annotated as compared to other cell types. We implemented both “1 vs 1” and “1 vs all” logistic regression to identify marker genes in each cluster as well as between two clusters in this study. The datasets used in this study are:

### GSE109999: A Gold-Standard Immune Cell Dataset^32^

This dataset consisted of FACS-sorted single cells representing B-cells, granulocytes, erythroblasts, and progenitor cells sorted from bone marrow, and T-cells isolated from the Thymus of 10-13 weeks old female C57/BL6 mice. These isolated cells were subsequently pooled together and subjected to sequencing using the CEL-seq2 protocol. Since the biological cell types were FACS-sorted prior to sequencing, this dataset can be labeled as a gold-standard reference for cell type identity.

### Tabula Muris

Tabula Muris, a multifaceted compendium of single-cell transcriptome data derived from the model organism *Mus musculus*, comprises nearly 100,000 cells hailing from 20 distinct organs and tissues^31^. In our study, we used a subset of the Tabula Muris dataset, specifically originating from the bone marrow. The Tabula Muris dataset comprises two methods for transcriptomic analysis: One utilizes microfluidic droplet-based 3’-end counting, which surveys thousands of cells per organ with relatively low coverage (Droplet). The other employs FACS-based full-length transcript analysis, providing higher sensitivity and coverage for more detailed insights from fewer cells (FACS).

### Human Datasets

To assess the stability of scSCOPE and other DGE methods, we used PBMC human datasets generated by Ding et. al in the same tissue by using multiple methods, including Dropseq, 10x V2, 10x V3, Dropseq, inDrops, Cel-Seq and Seqwell^30^.

### Marker Gene Identification using other methods

Differential expression tests for Wilcox Rank Sum’s test, MAST, DESeq2, Bimod and ROC were carried out using the FindMarkers() function from Seurat^20^. An average log fold change cut-off of 0.5 was used and the marker genes not expressed by at least 45% of cells in any group were discarded.

## Data Availability

Publicly available scRNA-seq datasets for methods benchmarking: Mouse Immune (GSE109999), Human PBMC (GSE132044), Tabula Muris (https://figshare.com/projects/Tabula_Muris_Transcriptomic_characterization_of_20_org ans_and_tissues_from_Mus_musculus_at_single_cell_resolution/27733).

## Code Availability

scSCOPE is freely available to all users from our GitHub website (https://github.com/QingrunZhangLab/scSCOPE).

## Acknowledgements

J.G. is a New York Stem Cell Foundation Robertson Investigator. This research was supported by The New York Stem Cell Foundation (NYSCF-R-N163 to J.G.), the Canadian Institutes of Health Research (RN418797-438409 to J.G.) and the National Science and Engineering Research Council (RGPIN-2019-04820 to J.G.). We thank the support from the HBI International Graduate Recruitment in Neuroscience Scholarship (S.A.) and ACHRI Graduate Scholarship (S.A.). We would also like to thank Mehr Malhotra for proof-reading our article.

## Competing interests

The authors declare no competing interests.

Supplementary Table 1: Number of marker genes and pathways identified for each cluster by different methods in gsBlood and Human Datasets.

Supplementary Table 2: Marker Genes Identified for all datasets used in this study by scSCOPE.

Supplementary Table 3: Pathways identified and ranked for gsBlood and Tabula Muris (B-cells) datasets by scSCOPE.

Supplementary Table 4: Number of genes identified by scSCOPE and other different methods in different pathways enriched in separate comparisons in the gsBlood dataset.

---

**Supplementary Fig. 1:**
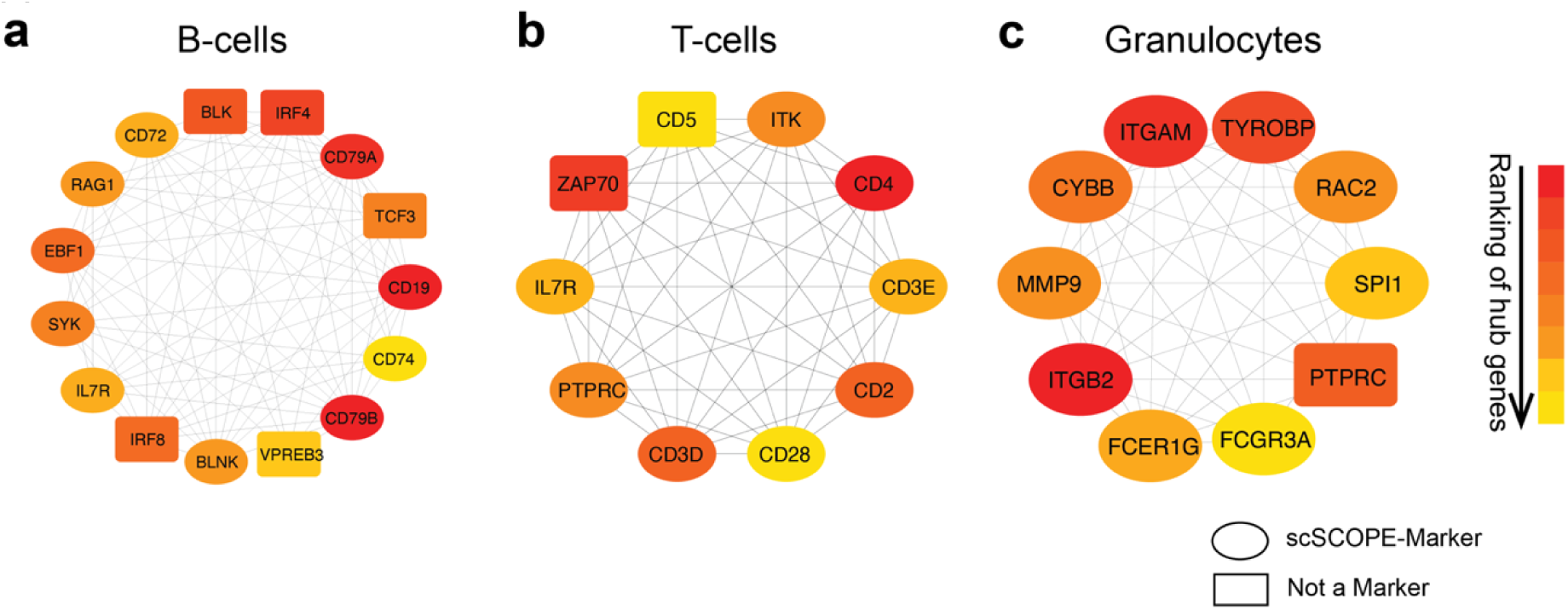
Correlation of Hub Genes and scSCOPE identified marker genes in gsBlood Dataset. Top Hub Genes identified in **a** B-cells **b** T-cells **c** Granulocytes of gsBlood dataset are shown as examples. scSCOPE identified markers are indicated by oval shape. Hub genes are ranked based on their level of gene co-expression, indicated by a red-yellow color theme.

**Supplementary Fig. 2:**
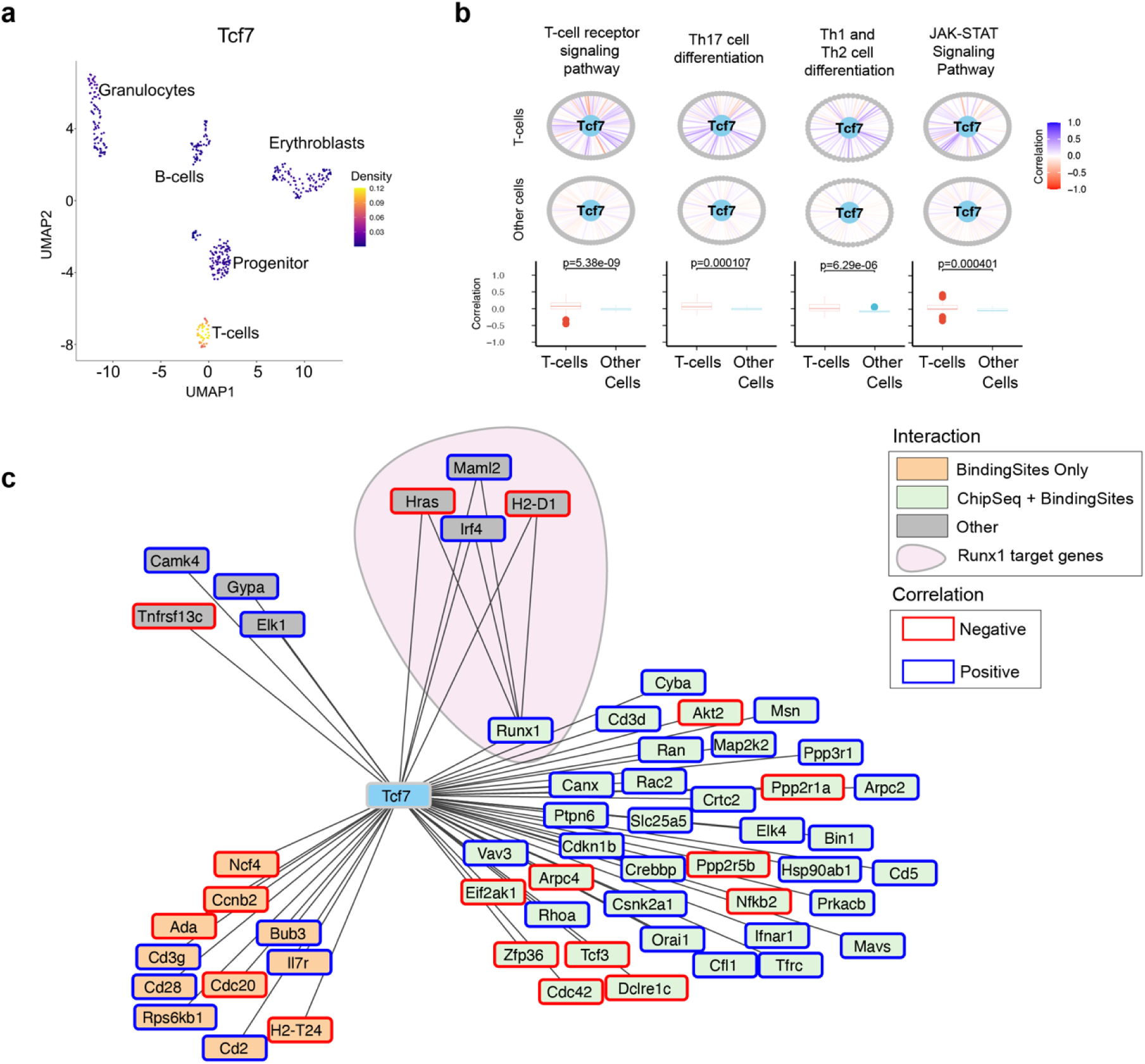
scSCOPE reveals potential targets of *Tcf7* gene in T-cells. **a** Density Plot showing the relative expression of *Tcf7* gene in different cell types of gsBlood dataset. **b** Gene Network Plots for *Tcf7* gene show the extensive interactions of *Tcf7* gene with genes across multiple pathways in T-cells. In each pathway, *Tcf7* gene is placed in the middle with all other genes in the pathway placed in the circumference of the circle. The lines connecting *Tcf7* to these genes indicate Pearson’s correlation coefficient, ranging from −1 (red) to +1 (blue), reflecting the strength and direction of correlation. Correlations are separately calculated for two distinct groups: in this case T- cells and all other cell types. Boxplots accompanying the plots contrast the distributions of correlations between these groups. Additionally, p-values from the Kolmogorov- Smirnov test are provided to assess the statistical significance, with the null hypothesis stating that two samples are drawn from the same distribution. **c** Network diagram showing the co-expressed genes of *Tcf7* identified by scSCOPE and their annotations based on published experimental results.

**Supplementary Fig. 3:**
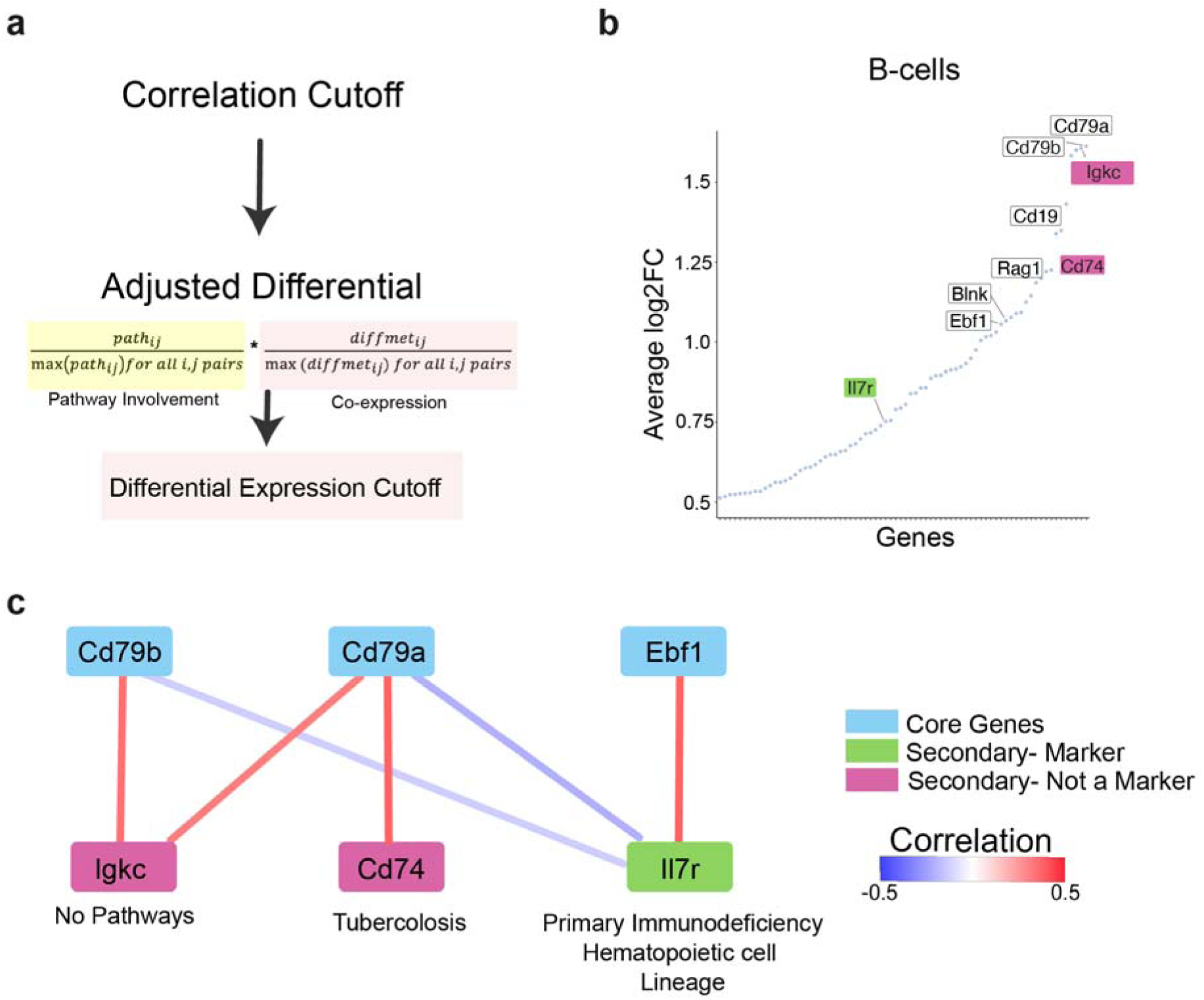
scSCOPE can identify genes with low fold change but extensive interactions as marker genes. **a** Genes identified as core and secondary by scSCOPE must pass correlation cutoffs, adjusted differential cutoffs, and differential expression cutoffs to be classified as marker genes. **b** Scatter Plot showing the average log2FC of DEGs identified by Wilcox rank sum method for B-cells in gsBlood dataset. Top marker genes for each cluster are labelled inside a box, indicating their ranking among DEGs from the Wilcoxon analysis. Although *Il7r* ranks low in terms of average log2FC in B- cells, scSCOPE identifies it as a marker gene in B-cells. **c** The *Il7r* gene is co-expressed with three core genes identified for B-cells and is also involved in two B-cell pathways.

**Supplementary Fig. 4:**
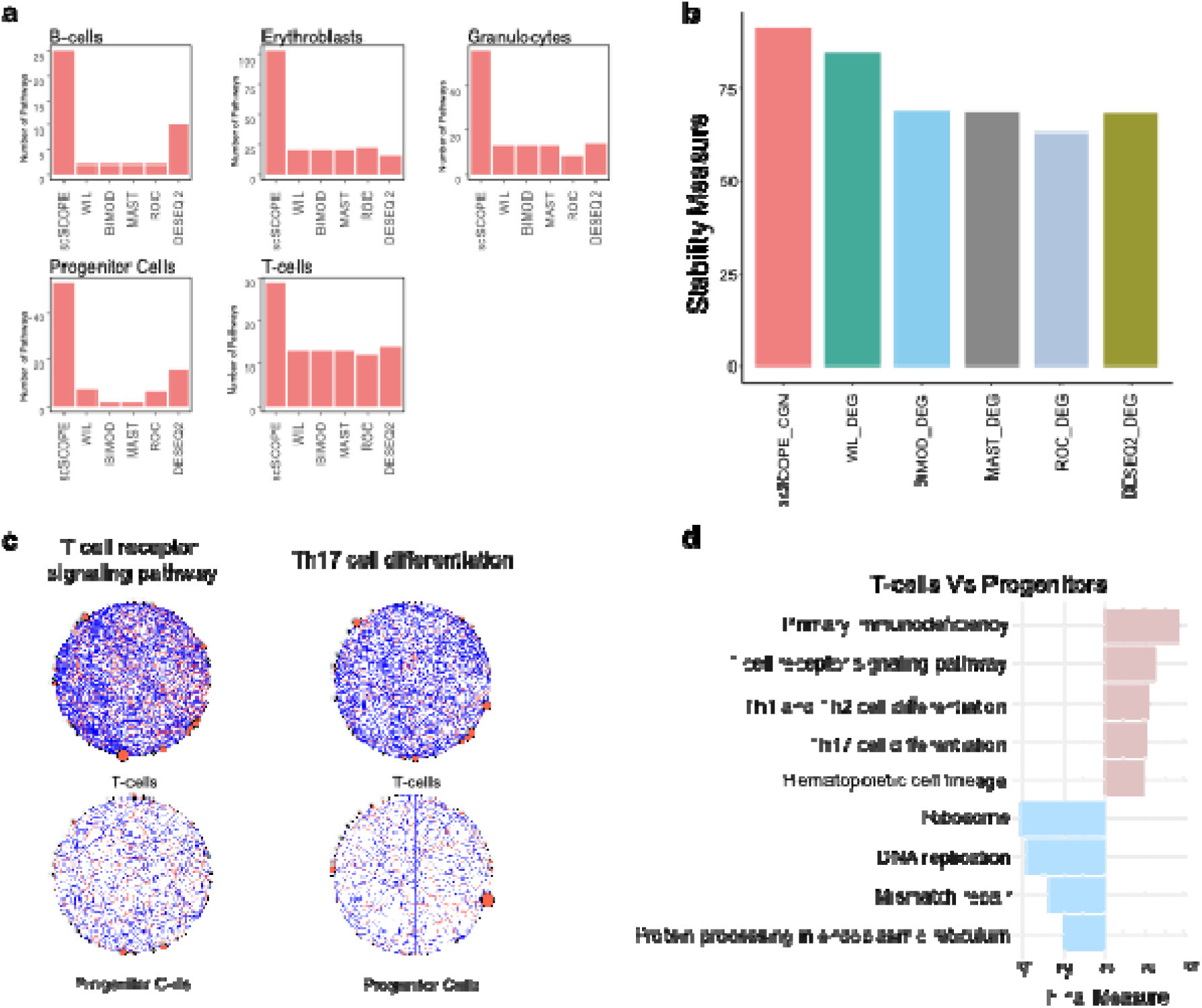
scSCOPE identifies a higher number of and more stable pathways. **a** Bar plot shows that scSCOPE identifies more pathways across all clusters as compared to other methods in gsBlood dataset. ORA was used to identify pathways for DEG methods. **b** The stability of each method in identifying pathways in the same cluster across different human PBMC datasets was measured. **c** Pathway Network Plots show the difference in expression and co-expression patterns of all the genes within T-cell Receptor Signaling Pathway and Th17 cell differentiation pathway between T-cells and Progenitor Cells. Pathway Network Plots are constructed for both T-cells and progenitor cells, facilitating a comparative analysis of pathway dynamics between the two cell types. In each plot, all the genes in the pathway are placed in the periphery of the circle. Genes are colored as orange (marker genes identified by scSCOPE) or gray. The size of each node corresponds to the average expression of the gene in the group, while edges connecting the nodes represent Pearson’s Correlation between two genes, with thickness indicative of correlation strength. Blue edges signify positive correlations, while red edges indicate negative correlations. **d** Pathway Bar Plot revealing the top pathways identified for T-cells versus progenitor cells, utilizing the novel metric “corrExpress.” Both “posCorrExpress” and “negCorrExpress” are combined to be named as “Final Measure”. This metric integrates differences in both gene-expression and gene-gene co-expression across all genes within the pathway. Pathways depicted with baby pink bars predominantly feature upregulated genes in T-cells, while those with light blue bars denote an abundance of upregulated genes in progenitor cells.

**Supplementary Fig. 5:**
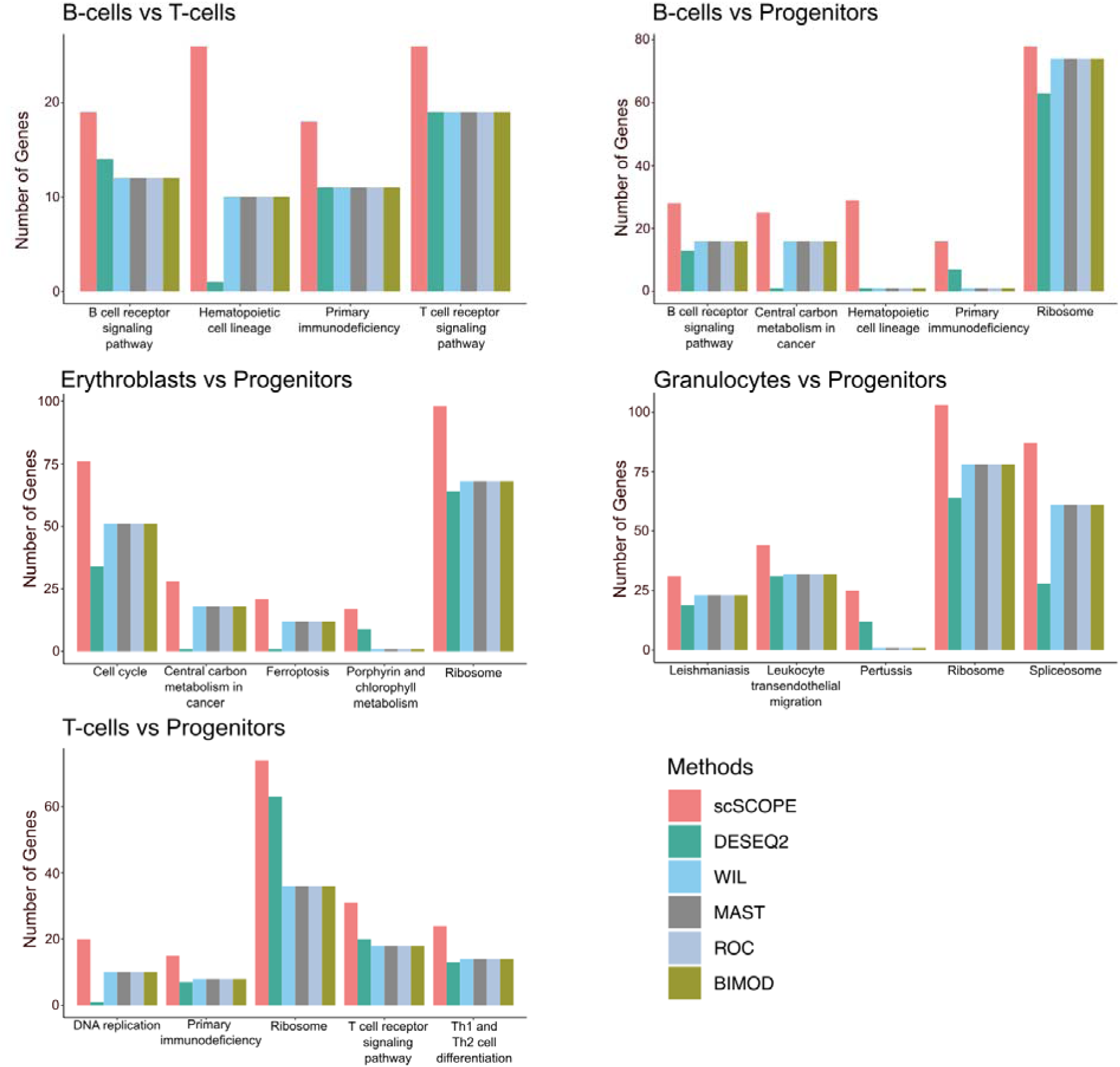
scSCOPE identified more genes within each pathway as compared to other methods. Bar Plots show the number of genes identified by various methods enriched in different pathways across different clusters in gsBlood dataset. The pathways were chosen from the top pathways identified by scSCOPE for each comparison (Supplementary Table 3). For every pathway in all clusters, scSCOPE identifies higher number of enriched genes than other methods.

## Notes

### Competing Interest Statement

The authors have declared no competing interest.

